# Mesenchymal stromal cell encapsulation in uniform chitosan beads using microchannel emulsification

**DOI:** 10.1101/2022.08.18.500253

**Authors:** Dongjin S Shin, Francesco K Touani, Damon G.K Aboud, Anne-Marie Kietzig, Sophie Lerouge, Corinne A Hoesli

## Abstract

Mesenchymal stromal/stem cells hold potential in repairing damaged tissue through paracrine effects. Their delivery though injectable biodegradable microbeads can improve cell retention and survival at the infusion site. A stirred emulsion process was previously implemented to immobilize these cells in injectable chitosan microbeads for cell therapy applications, but this process leads to broad bead size distribution (coefficient of variation > 40 %). Polydisperse beads may negatively affect the viability of the entrapped cells through oxygen limitations, damage to larger beads during injection, and reduced control over the cell payload and treatment reproducibility. The objective of this work was to modify a microchannel emulsification system initially designed for alginate-based encapsulation to immobilize mesenchymal stromal/stem cells in monodisperse chitosan microbeads. The main factors (e.g., microchannel geometry, chitosan solution viscosity, interfacial tension and flow rate) affecting droplet generation and diameter were investigated. The adapted process enabled the production of monodisperse chitosan microbeads with controlled sizes ranging from 600 µm – 1500 µm in diameter at a coefficient of variation less than 10 %. In a single pass through a 21 G syringe needle (ID: 513 µm), the fraction of ruptured beads was significantly reduced for microchannel-generated vs stirred emulsion-generated beads with matching volume-weighed bead diameter (D[4,3]). The viability of the immobilized cells immediately after the process was 95 % ± 2 % and no significant difference in cell survival and growth factor secretion was observed between microchannel and stirred emulsion-generated beads over 3 days of culture. Future directions include channel multiplexing to increase throughput for clinical applications. Although the device was developed for cell encapsulation, this process could be implemented for encapsulation of other biomolecules, bioactive or living cell agents for applications in the food and drug industry.

## INTRODUCTION

Due to their multipotency and secretion of pro-angiogenic and immunosuppressive factors, mesenchymal stromal/stem cells (MSCs) show great potential for repairing endogenous tissues, a process that dramatically deteriorates with age ^1–3^. Although implanting MSCs is an appealing strategy for tissue regeneration, cell death shortly after administration was shown to reduce therapeutic effects ^4^. Additionally, low cell retention and engraftment prior to vascularization have limited the successful application of MSCs in regenerative medicine ^5^.

Instead of injecting free-floating cells, a potential alternative is to immobilize cells in either micro- or macroscale hydrogels that allow bidirectional diffusion of nutrients, oxygen, and cell signalling factors ^6–8^. Macroencapsulation immobilizes cells in a single encapsulation matrix, whereas microencapsulation entraps the cells in many microspheres ranging from a few micrometers to millimeters in diameter ^9, 10^. The higher surface area/volume ratio of microbeads compared to macrogels can improve diffusion-based oxygen delivery to the immobilized cells to reduce hypoxic cell death ^8^. This can also favour integration of the therapeutic cells into the surrounding environment as the biomaterial degrades ^6^.

Chitosan, a cytocompatible and biodegradable amino-polysaccharide obtained by deacetylation of chitin ^11^, is a particularly interesting candidate for cell encapsulation, due to its tunable biodegradation rate. While most solubilization methods require acidification which is problematic for cell processing ^12^, the issue can be overcame by adding a weak base such as beta-glycerophosphate to the solubilized chitosan solution prior to mixing with cells. This leads to a solution with neutral pH at room temperature which undergoes sol-gel transition at body temperature ^13^. The formulations that we have developed with a mixture of hydrogen carbonate and phosphate enabled to form cell-loaded injectable chitosan hydrogels with good mechanical properties and rapid gelation of the chitosan solution under physiological conditions (pH 7-8, 37 °C) ^14, 15^.

In previous work, we successfully generated chitosan microbeads via stirred emulsification ^16^. The method, originally described by Poncelet et al. for hydrogels undergoing internal gelation^17^ and adapted to alginate-based cell encapsulation by Hoesli *et al*. ^18–20^, generates emulsion droplets using a rotating impeller in a pre-heated oil bath, allowing rapid gelation. Although this method was suitable to produce injectable chitosan hydrogels, due to non-uniform energy dissipation in stirred systems, the resulting beads were polydisperse ^16^. Since their broad size distribution may lead to heterogeneity in cell survival and make it difficult to control the number of cells injected, methods that produce beads of uniform diameter at high throughput are of interest in the cell encapsulation field ^21, 22^.

The goal of this project was to develop a method to achieve small and narrow size distribution chitosan microbeads for MSC encapsulation. We modified a microchannel emulsification (MCE) device previously designed by Bitar and Markwick et al. to encapsulate pancreatic ß cells in alginate hydrogel ^23^. Although the bead diameter was relatively large (beads ranging from 1.5 mm – 3 mm in diameter), the size distribution was narrower (coefficient of variation < 10 %) than the stirred emulsification method (coefficient of variation > 60%). Here, we adapted the microchannel system for chitosan-based MSC encapsulation and evaluate the suitability of the new method for cell therapy applications. By modifying the device structure and optimizing system variables, we generated monodisperse chitosan microbeads while maintaining high cell viability.

## MATERIALS & METHODS

### Hydrogel preparation

Shrimp shell chitosan powder (ChitoClear, HQG110, *M*_*w*_: 150-250kDa, degree of deacetylation 93%, Primex Iceland) was solubilized in 0.12M HCl solution using an overhead stirrer at 900 rpm for three hours. This solution was sterilized by autoclaving for 20 min at 121°C, 15 psi. The gelling agent solution was a mixture of two weak bases: sodium hydrogen carbonate *NaHCO*_3_(G9422, Sigma-Aldrich) and phosphate buffer. ^15^ Phosphate buffer solution was prepared at 0.1 M (pH 8.0) by solubilizing sodium phosphate dibasic *Na*_2_*HPO*_4_(S5136, Sigma-Aldrich) and sodium phosphate monobasic *NaH*_2_*PO*_4_(S2554, Sigma-Aldrich) in Milli-Q water at 2.65 × 10^*−*2^ g/mL and 0.16 × 10^−2^ g/mL, respectively. Then, sodium hydrogen carbonate was solubilized in phosphate buffer solution at 0.375M. The pH of the gelling gents was measured using a pH meter (LAQUAtwin pH-22, Horiba). The gelling agent solution (phosphate buffer 0.1M; sodium hydrogen carbonate 0.375M) was passed through a 0.22 µm filter (Corning) for sterilization ^16^. Chitosan solution and gelling agent were mixed at a 3:1 volumetric ratio using syringes and a Female Luer lock connector to prepare for the aqueous phase. This solution was mixed with the cell stock (for experiments with cells) or serum-free (for experiments without cells) at a 4:1 volumetric ratio, leading to a chitosan precursor solution (phosphate buffer 0.02M; sodium hydrogen carbonate 0.075M) containing 0.5 × 10^6^ cells/mL. The serum-free medium used for encapsulation was alpha minimum essential medium (alpha-MEM, Gibco) supplemented with 10 % bovine serum albumin (BSA, A4812, Sigma-Aldrich).

### Cell culture/Cell stock preparation

Bone marrow-derived human MSCs (Lonza Inc.) were cultured using NutriStem XF (biological Industries) medium supplemented with 0.6 % MSC NutriStem XF Supplement up to 90 % of confluence based on microscope observation. Cells were seeded at 2860 cell/cm^2^ density on T175 tissue culture flasks (Canted Neck Red Ventilated Cap for Adherent Cells, 50-809-259, SARSTEDT Inc.) with medium volumes of 0.142 mL/cm^2^. Immediately before encapsulation, the cells were detached with 0.03 mL/cm^2^ of Trypsin/EDTA (Wisent) and incubated for 3 - 5 min. Then, the cell suspension was transferred into a 50 mL centrifuge tubes and centrifuged at 500 x g for 5 min. The cells were re-suspended in serum-free medium at 4-fold the desired encapsulated cell concentration.

### Interfacial tension measurement

The interfacial tension between the oil phase and the chitosan was measured at room temperature by the pendant drop method using a goniometer (Future Digital Scientific Corp.) connected to a video camera system and computer software (SCA20, Dataphysics). A quartz cuvette was filled with chitosan solution. A 500 µL syringe needle (523159, Hamilton) filled with oil was immersed into the solvent and a droplet was created by ejection at 36 µL/s. The drop profile of the oil phase suspended in the chitosan solution was recorded using a high-speed camera. The interfacial tension was estimated by fitting the Young-Laplace equation to experimental images using OpendropV1.1 software (Supporting information) ^24^. For the fitting process, the differences in density of the liquids were considered (chitosan mixture density of 1.1 g/cm^3^; Novec oil density of 1.614 g/cm^3^). For each sample solution, the interfacial tension value was the average of three measurements.

### Viscosity measurement

The apparent viscosity of the chitosan precursor solution was measured at room temperature (~22 °C) using a MCR302 Rheometer (Anton Paar) with a cone-plate system (strain and frequency at 1% and 1 Hz, respectively) immediately after mixing. Shear rate was ramped logarithmically from 0.1 s^−1^ to 200 s^−1^ for a total of 15 s. The inspection time was 1 s and the duration of inspection value was 0.5 s for each data point. Milli-Q water was used for the precursor solution preparation instead of gelling agent to avoid viscosity changes during the measurement.

### Microchannel device and channel fabrication

The device consists of a microchannel plate sandwiched between a bottom and top chamber designed to be easily assembled/disassembled (Supporting information 1). The top and bottom chambers were designed (Supporting Information) using AutoCAD (AutoDesk©) software and machined from acrylic rods (8528K55, Clear Scratch- and UV-Resistant Cast Acrylic Rod, 4 - 1/2” Diameter, McMaster-Carr). Microchannels were machined from polytetrafluoroethylene (PTFE) sheets (9266K11, Chemical-resistant slippery PTFE Sheets, McMaster-Carr) of either 1.635 mm or 0.86 mm in thickness via femtosecond laser micromachining. The laser setup consists of a Libra Ti: Sapphire laser system (Coherent Inc.) with a central wavelength of 800 nm, pulse duration < 100 fs, and a 1 kHz repetition rate. The pulse energy used was 200 µJ, and the beam was focused using a 200 mm focal length convex lens to a spot radius of 20 µm, leading to a pulse peak fluence of 30.7 J/cm^2^. The surface was micromachined in a raster scanning pattern using a translation speed of 2 mm/s and a line spacing of 20 µm. High resolution 3D images of the channels were acquired using a laser confocal microscope (Olympus OLS 4000 LEXT) to measure the resultant channel dimensions.

### Dynamic contact angle measurement

Water contact angles were measured by dispensing 2 μL reverse osmosis water droplets on test surfaces using a 32G syringe needle (Hamilton). Then, the liquid volume was increased to 7 μL at a rate of 0.25 μL/s. After a five second pause, all the liquid was drawn back into the needle. The advancing and receding contact angles of the resulting videos were measured using SCA20 software (DataPhysics Instruments, USA).

### Scanning electron microscopy

PTFE surfaces were sputter-coated with platinum to develop a 5 nm thick conductive layer on the surface using an EM ACE600 sputter coater (Leica). Images were acquired using a Quanta 450 scanning electron microscope (FEI company).

### Microbead production by microchannel emulsification

The bottom chamber inlet port was connected (5117K85, McMaster-Carr) to 5 cm length Tygon tubing (Saint-Gobain) with a female Luer to hose barb connector at the other extremity. Two O-rings (ID: 3 mm OD: 7 mm, 5233T474, McMaster-Carr) were placed in O-ring grooves located (a) between the microchannel plate and the top chamber and (b) the microchannel plate and the bottom chamber to create a water-tight seal (Supporting information 1). When aseptic device assembly was required (i.e. for cell culture experiments), these device components were sterilized through overnight immersion in a 70 % ethanol bath and air-drying for 30 min in a biosafety cabinet followed by aseptic handling. The microchannel plate was placed between the top chamber and the bottom chamber and assembled using four tightening screws (**Figure 1.A**). The chitosan precursor solution was prepared in a syringe with the desired volume (3 mL - 5 mL, Beckton-Dickinson) as explained and mounted onto the syringe pump (Sage Instruments mode 355/365, Cole-Parmer) (**Figure 1.B**). To prepare the oil phase for microchannel emulsification, Fluoro-Surfactant (RAN Biotechnologies Inc.) was dissolved in 3M™ Novec™ 7500 (3M Company) at 0.066% w/w and aseptically filtered (0.2 µm syringe filters, SARSTEDT). A total of 70 mL of oil phase was prepared per batch. Out of 70 mL, 50 mL was preheated to 37 °C, and the rest (20 mL) was stored at 22 °C. Then, the aqueous phase was infused into the bottom chamber until the solution level reached the microchannel plate (1 mL ~ 1.5 mL). To promote rapid gelation at the oil surface but avoid early gelation at the interface, 20 mL of the 22 °C oil was poured into the top chamber followed by gradual addition of 50 mL of the 37 °C oil (Figure 1.B). While the preheated oil was slowly being added, the aqueous phase was pumped into the bottom chamber, initially with a flow rate of 1 mL/min to purge air trapped inside the channel, which was steadily decreased to the desired flow rate for droplet production. The droplets detached from the microchannel floated to the oil surface. When the desired volume of droplets was obtained at the collection site, the top was air-tight sealed with a sterilized parafilm and the entire device was placed in an incubator to ensure complete gelation at 37°C for an additional 5 min - 10 min. Then, the device was taken out of the incubator and 10 mL of wash solution (10mM of HEPES and 170 mM NaCl, pH 7.4, 10% v/v bovine serum albumin) was added into the batch. The fully gelled chitosan microbeads were transferred to a 50 mL centrifuge tube using a spatula and centrifuged at 500 x g for 1 min to separate Novec oil from HEPES buffer saline solution and the chitosan beads. The chitosan beads were transferred to a 100 μm nylon cell strainer (22363-549, Fisherbrand^tm^) and rinsed by evenly dispensing 5mL of wash solution over the beads. A volume of 200 µL of beads suspension were transferred in a 24-well plate (9023511, SARSTEDT) and 800 µL of low-serum medium (0.2% v/v FBS in alpha-MEM) were added in each well plate. Half of the beads were used for cell viability measurements immediately after the encapsulation and the other half were cultured in low-serum medium for 3 days.

**Figure 1.**
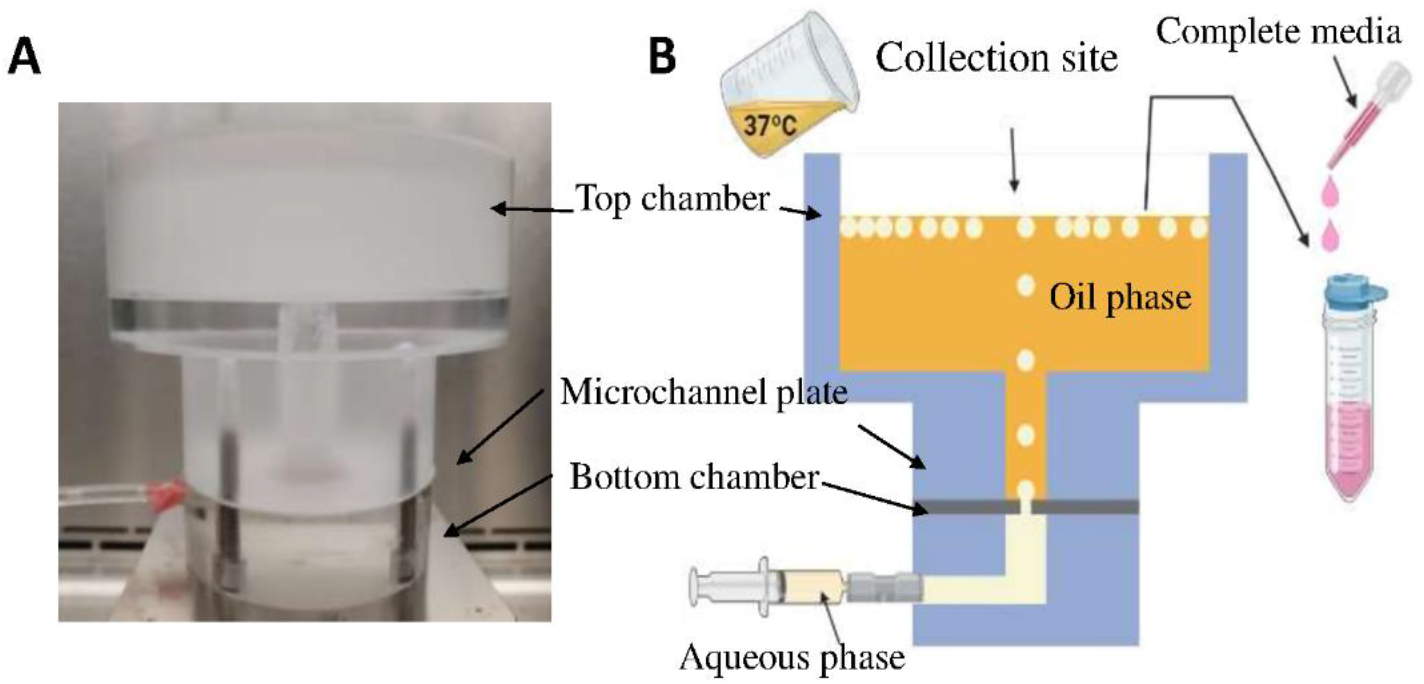
Microchannel emulsification device and the microbead production process. (**A**) photograph of the microchannel device (**B**) Schematic drawing of the encapsulation method using the microchannel emulsification device.

### Microbead production by stirred emulsification

Chitosan microbeads were generated via stirred emulsification as previously described ^16^. Briefly, 20 mL of light mineral oil (O121-4, Fisher) was placed in a 100 mL microcarrier spinner flask (Bellco Glass Inc.) located in a preheated in water bath (37 °C). The water bath was placed on a magnetic stirrer (VWR), temperature set at 37°C. With continual agitation at 600 rpm, the aqueous phase was prepared as mentioned above and injected into the preheated oil through a 18G syringe needle (1’’ in length, Air-Tite) drop by drop immediately after chitosan preparation. At the 3-min mark, the rotational speed was gradually decreased to 200 rpm and beads were left to gel for 7 min at this mixing speed. At the 10-min mark, 20 mL of wash solution was added. The contents were transferred to 50 mL centrifuge tube and centrifuged at 500 x g for 1 min to separate mineral oil from HEPES buffer saline solution and the chitosan beads. After removing the oil, the beads were resuspended and transferred to a 100 μm nylon cell strainer for washing. Then, beads were transferred and stored in a 24-well plate using the method described for microchannel emulsification. The impeller rotational speed (600 rpm) was determined to match the D[4,3] (De Brouckere mean diameter, also called volume-weighted mean diameter) of the microchannel emulsification beads’ diameter.

### Microbead size and size distribution

Beads (0.5 mL) were placed in 10 mL of 0.01 g/L Eosin Y (Fisher Chemical^tm^) in wash solution. The solution was stirred for 20 min on a rotary shaker. Images of stained beads dispersed in wash solution were acquired using an UVP Biospectrum Imaging system. The diameter equivalent to the surface area in 2D images was determined using ImageJ freeware (Fuji) ^25^. The D[4,3] ^18, 26^ and coefficient of varication (C.V.) were calculated from the diameter distributions of each batch using JMP (SAS) software.

### ead mechanical properties

To evaluate microbead mechanical properties and viscoelasticity, we used a MicroSquisher (CellScale) with a parallel-plate compression configuration as described in the previous work ^16^, using a 305 µm diameter microbeam. For the measurement, hand-picked microbeads of 700 µm – 800 µm diameter were placed onto a metal platform immersed in HEPES buffered saline solution at room temperature. Each bead went through three consecutive cycles of up to 30% volume compression (30 s loading time, 10 s hold time, and 30 s recovery time) to measure the force as a function of displacement using the accompanying software (SquisherJoy). The compressive modulus (E) of each microbead was calculated using the modified Hertzian half-space contact model (Supporting information 2) ^27^.

### Injectability

Cell-free microbeads of matching matching D[4,3] were generated using microchannel and stirred emulsification. Microbeads (1 mL) were suspended in wash solution at different dilutions (3 mL and 6 mL). Bead suspensions (total of 200 µm) were pipetted and transferred to a syringe (1mL) and the beads were manually (to reproduce handling for animal or clinical studies) injected through 21 G (inner diameters of 513 µm) in 24 wells plate filled with 500 µL wash solution. Images were taken using phase-contrast light microscopy (Leica DM LB2) to quantify the number of ruptured or damaged beads.

### Viability of immobilized MSCs

Each sample of 200 µL bead suspension was transferred to a well (24-well plates) followed by addition of 800 µL serum-free alpha-MEM containing 2 µM Calcein AM and 5.5 µM ethidium homodimer. After 45 min incubation, live/dead images were taken randomly at five different locations per sample with an inverted fluorescence microscope (Leica DMIRB) at 50x magnification. The images were analyzed using Image J software (Fiji).

### Paracrine activity

The paracrine activity of the cells was assessed by measuring the concentration of vascular endothelial growth factor A (VEGF) released in the conditioned media (0.2% v/v FBS in alpha-MEM) of microencapsulated MSC (500 K/mL) after 3 days of incubation. Conditioned media samples were centrifuged (5 min, 300 x g), then diluted 1/3 in fresh conditioned media. Determination of the amount of VEGF was performed using the Quantikine ELISA Human VEGF Immunoassay (Biotechne).

### Statistics

Normality tests were conducted. For data with normal distribution, two-way comparisons were performed using t-tests. Multiple comparisons used two-way ANOVA (GraphPad Prism 9.4.0) followed by Tukey post-hoc tests. For non-normal distributions, two-way comparisons were performed using Mann-Whitney tests. The level of significance was determined based on p-value (ns = p > 0.05, *: p< 0.05). Results with replicates were represented as average ± standard error of the mean (GraphPad Prism 9.4.0). N represents the number of replicates and n represents the number of samples in each replicate batch.

## RESULTS AND DISCUSSION

The main challenges in encapsulating MSCs in chitosan beads via microchannel emulsification were to assure sufficiently rapid bead gelation to avoid coalescence in the collection bath, and to obtain microbeads that are small enough to be injectable. To achieve this objective, an iterative design process was required to optimize the device structure, microchannel geometries, and surfactant addition.

### Microchannel dimensions and surface properties

To determine the effect of channel geometry on bead size and formability, we compared straight-through symmetric microslot channels (**Figure 2A**) and asymmetric channels (combination of microslot and cylinder orifice, **Figure 2B)** achieved by varying the channel width, height, and slot depth (**Table 1)**. We kept the aspect ratio of the outlet length/width to be over 5 assuming that a minimum aspect ratio is required to establish a unidirectional laminar flow ^28^. Due to limitations in laser micro-machining, the walls of the ablated microchannels have a tapering angle ranging from 83 to 86⁰ as the laser mills deeper into the plate, resulting in dimensional differences between the microslot inlet and the outlet widths. This occurs as a result of the shape of the focused laser beam, which forms a cone after passing through the focusing lens (**Table 1)**. Contrary to the microslots, the cylinder orifice showed minimal dimensional tapering. This might be because of different laser-milling strategies used for creating microslots and cylinder orifices. In contrast to the raster scan trajectory used for the microslots, the laser beam spiraled in a circular shape to generate cylinder orifices. Due to the limitation of the focused laser beam’s spot size, the minimum cylinder diameter achieved was 220 µm.

**Figure 2.**
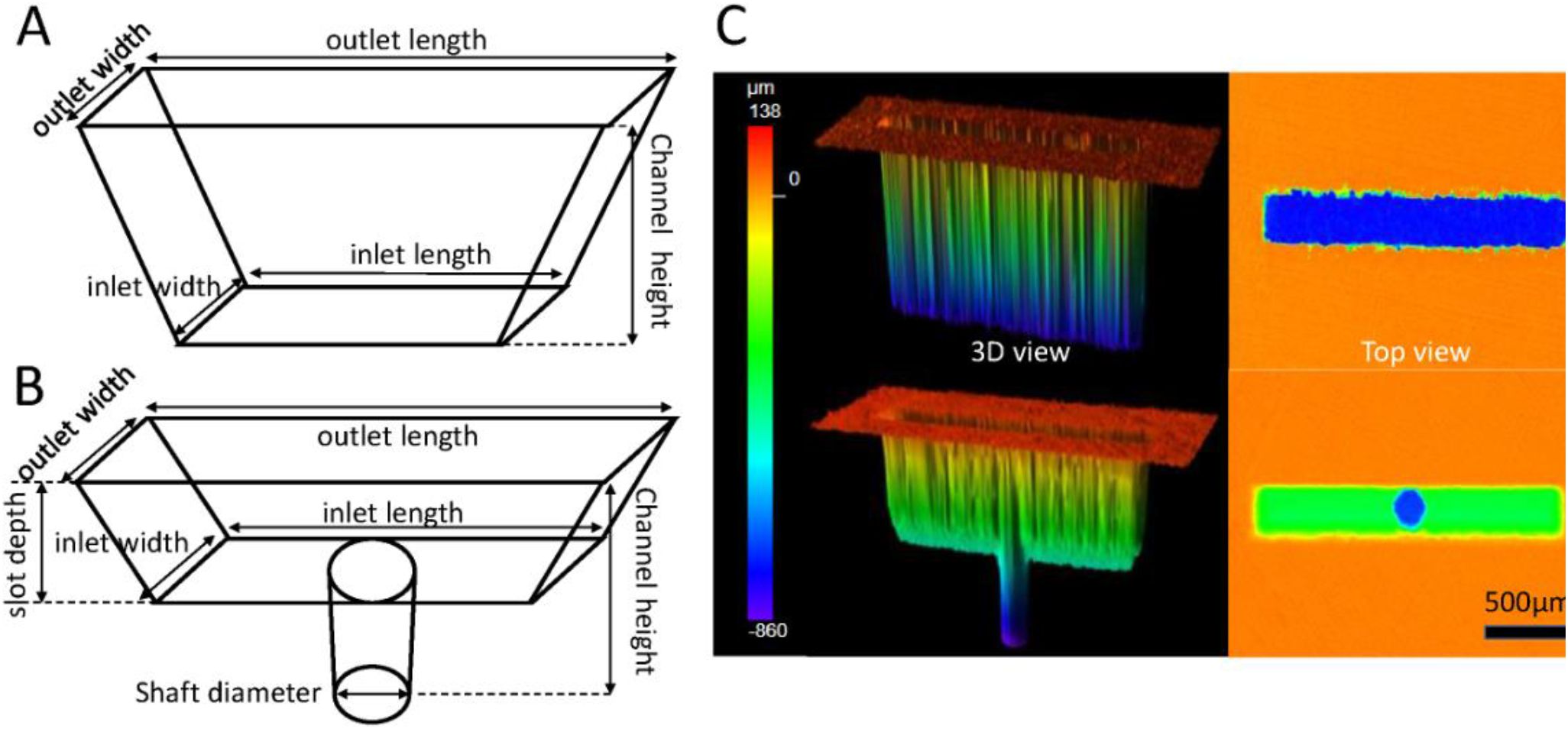
Channel geometry. (**A**) Three-dimensional drawing of a symmetric channel (**B**) Three-dimensional drawing of an asymmetric channel. The length difference between the outlet and the inlet was exaggerated in the drawings to emphasize dimensional tapering. (**C**) Confocal heatmaps of two different channels.

**Table 1.** Dimensions and labels of channels used for the study. For most microchannels, only one channel was manufactured with the exception of **Asym C** (N > 20).

| Channel type | Channel label | Rectangular duct width (outlet/inlet) ($\mu\text{m}$) | Cylinder diameter (inlet) ($\mu\text{m}$) | Hydraulic diameter (inlet) ($\mu\text{m}$) | Channel height ($\mu\text{m}$) |
| --- | --- | --- | --- | --- | --- |
| Symmetric | Sym A | Intended: 200/200<br>Result: 514/223 | N/A | 401 | 1635 |
|  | Sym B | Intended: 150/150<br>Result: 340/126 | N/A | 188 | 860 |
| Asymmetric | Asym A | Intended: 250/250<br>Result: 483/350 | 320 | 320 | 1635<br>(slot depth: 900) |
|  | Asym B | Intended: 200/200<br>Result: 358/242 | 238 | 238 | 1635<br>(slot depth: 900) |
| | Asym C | Intended: 150/150<br>Result: $300 \pm 11/220 \pm 10$ | $210 \pm 10$ | $210 \pm 10$ | 860<br>(slot depth: $500 \pm 10$) |
|  | Asym D | Intended: 150/150<br>Result: 315/235 | 220 | 220 | 860<br>(slot depth: 275) |

### Hydrophobicity of the microchannel

The microchannel surface should be as hydrophobic as possible to maximize oil phase wetting to promote droplet formation ^23, 29^. As expected ^30^, the hydrophobic pristine PTFE plates used to fabricate the had water contact angles above 90° (advancing water contact angle of *θ*_*A*_ = 97 ± 1°; receding water contact angle of *θ*_*R*_ = 43 ± 12°). The laser irradiation process results in a hierarchical topography consisting of a rippled microstructure that is superimposed by porous, wiry nanostructures (**Figure 3.A, B**). This porous structure is known to trap air pockets, causing water to wet the surface in the Cassie-Baxter wetting state, leading to superhydrophobic behaviour ^31, 32^. Using a surrogate surface with an irradiated area large enough to deposit water drops, we measured dynamic water contact angles of *θ*_*A*_ = 150 ± 4° and *θ*_*R*_ = 123 ± 8° (**Figure 3.C**). These large contact angle measurements confirm that the irradiated PTFE has low water wetting properties, which is expected to aid in droplet-pinch off.

**Figure 3.**
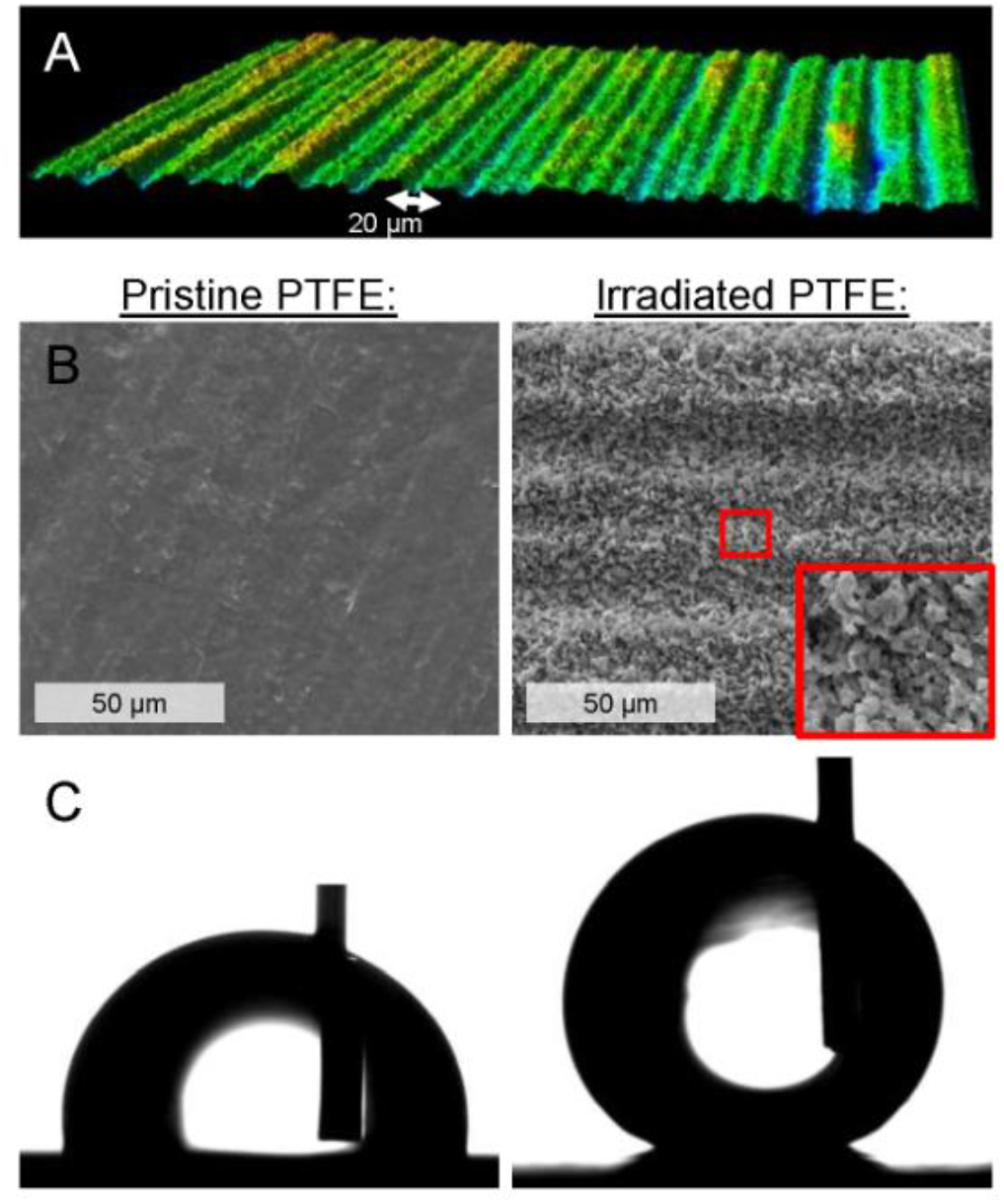
Superhydrophobic properties of channel surfaces obtained by laser micromilling. (**A**) confocal heatmap of the rippled microstructure. Ridges are spaced 20 µm apart. (**B**) Scanning electron micrographs of the pristine and laser irradiated PTFE. The red box in the irradiated image provides a zoomed view of the nanostructure. (**C**) Images of the advancing water contact angle on the pristine and irradiated material.

### Process development

We applied a temperature gradient to the oil phase to avoid sudden temperature increase in the aqueous phase at the plate surface which could cause premature chitosan gelation and channel obstruction. Among the gelling agent formulations known in previous work ^15^, we chose a formulation containing 0.02M phosphate and 0.075M sodium hydrogen carbonate based on its gelation kinetics (relative stability at room temperature and rapid gelation at 37°C, see supporting information 3). At higher phosphate concentrations, the precursor solution gelled too quickly, even at room temperature, resulting in generation of droplets with uncontrolled sizes as the overall processing time increases. At lower phosphate or carbonate concentrations, the gelation kinetics were too slow to obtain gelled beads in the collection reservoir.

Despite the temperature gradient established through this method, most droplets coalesced upon contact (**Figure 4A**). To stabilize the droplet interface and reduce coalescence during gelation, surfactants (*Fluoro-Surfactants*) compatible with water-in-fluorocarbon oil emulsions were added in the oil ^33^. The surfactant was effective in preventing bead coalescence at 0.066% w/w, as shown in **Figure 4B**. Also, *Fluoro-Surfactants* significantly reduced the interfacial tension between the two phases at this concentration in the pendant drop setting (oil-in-water), shown in **Figure 4C**.

**Figure 4.**
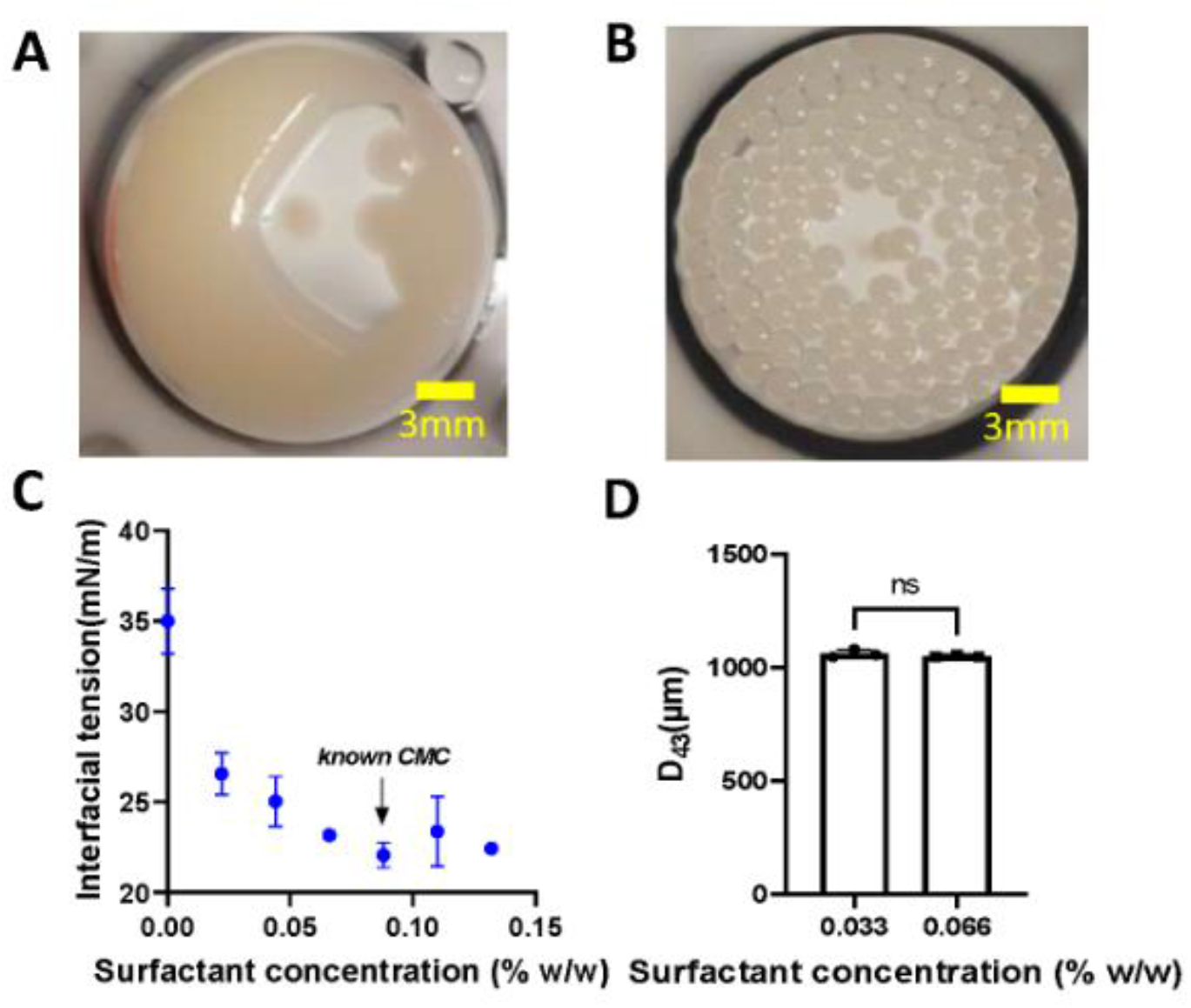
Effect of Fluoro-Surfactant on bead coalescence and the average bead size. (**A**) Coalesced droplets generated without the surfactants. (**B**) Droplets floating in the oil phase at 22C° with surfactant concentration of 0.066% w/w. Both images were taken 1 minute after the generation process. (**C**) Interfacial tension between the chitosan solution (1.6% w/v) and the oil phase at different surfactant concentrations (**D**) D43 of the beads generated at two different surfactant concentrations, critical micelle concentration (Critical micelle concentration, 0.066% w/w) and 0.033% w/w. (N=3, n > 750). Asym A (Table 1) was used for droplet generation.

As interfacial tension is one of the main driving forces in the droplet detachment process ^34^, we hypothesized that the surfactant concentration could potentially reduce the droplet size. However, the surfactant concentration had no significant effect on the average bead diameter (D[4,3]), as shown in **Figure 4D**. Unlike the pendant drop setting where droplet detachment takes more than seconds, the detachment time in the microchannel (milliseconds) may be too short for the surfactants to affect the interfacial tension ^35^. The rate of surfactant diffusion to the droplet interface may have been insufficient to impact droplet detachment while still creating a sufficient energy barrier to avoid later coalescence.

### Effect of channel geometry on the bead diameter

To reduce diffusion distances and enable eventual in vivo delivery via needle injection, bead diameters of reduced size (≤600 µm) were desirable. The beads generated using two different channel geometries with relatively similar inlet/outlet width and height (Sym A and Asym A) showed significant differences in average diameter (respectively, 2222 ± 31 µm and 1013 ± 10 µm, **Figure 5**). To determine whether a change in overall channel size could affect the bead diameter, we generated beads using channels with smaller inlet widths. For symmetric channels (Sym A vs. Sym B), a decrease in channel width showed no significant change in bead diameter (2222 ± 31 µm and 2105 ± 57 µm). As expected ^28^, reducing the asymmetric channel rectangular slot width (Asym A vs. Asym C) led to a significant reduction in bead diameter (1013 ± 10 µm and 573 ± 6 µm). The ratio of resultant bead diameter to the inlet width exceeded 10 in symmetric channels, whereas the ratio was 2-3 for asymmetric channels. Considering that the aspect ratio of the microslot is over 5, the result might indicate that asymmetric channels could effectively pinch off a droplet before the droplet diameter expands to reach the channel length ^35^, which may not have been the case for the symmetric channel under the conditions applied. Comparing channels of different slot depth (Sym B: 0, Asym C: 500 µm, Asym D: 275 µm) indicated that a minimum depth was required in asymmetric channels to achieve effective droplet pinch-off and significantly reduce the bead diameter.

**Figure 5.**
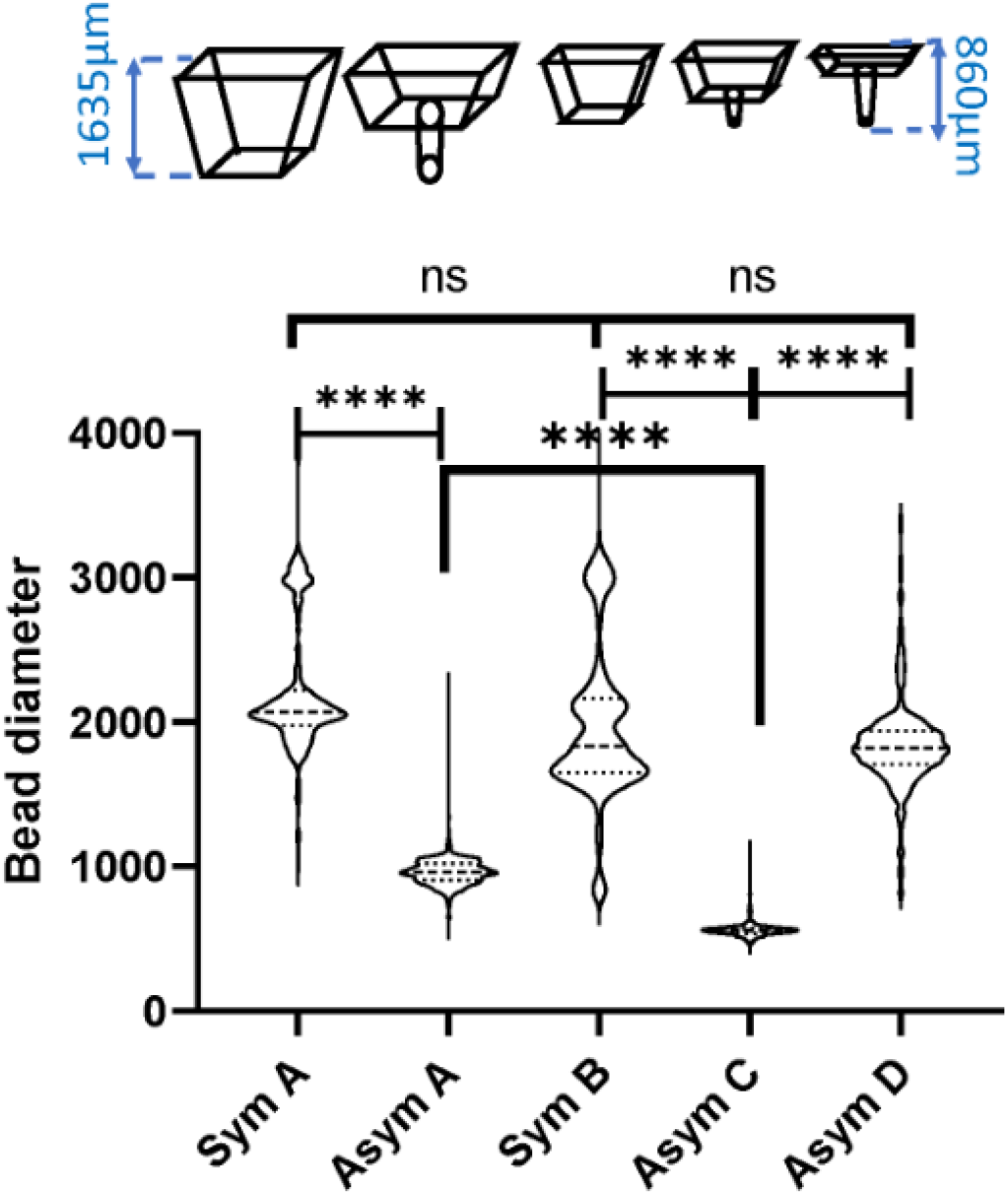
Effect of channel geometry on bead size distribution. Size distribution of beads generated by channels of (1) **different types** (Sym A and Asym A) and (2) different **slot depths** (Sym B, Asym C, Asym D). See detailed dimensions in **Table 1**. Each dotted line within the violin plot marks the quantile point dividing the range of the percentage distribution. (N > 2, n > 750, **** = p > 0.0001)

Larger (> 1500 µm) beads generated using symmetric channels tended to coalesce to form larger beads, likely due to incomplete gelation, resulting in bimodal or trimodal bead size distributions (C.V > 20 %). In contrast, the beads produced by asymmetric channels showed distinctive uninomodal size distribution (C.V < 10 %). From here on, to further decrease the size of the beads, we focused on optimizing the system using the channel with the smallest width (Asym C).

### Effect of aqueous phase viscosity on bead size

To study the effect of aqueous phase viscosity on bead diameter, we generated beads using precursor solutions with different chitosan concentrations at a fixed flow rate and quantified the bead diameter. To avoid the changes of viscosity with time, which occurs due to the reaction between chitosan polymers and gelling agent ^15^, we mixed chitosan with Milli-Q water instead of gelling agent for this part of the study. This measurement would represent a value near the initial pre-gelling viscosity of the chitosan mixture. The solution at the highest viscosity (2 % chitosan) generated broad bead size distributions. The dispersed solution often obstructed the channel leading to unstable flow observed as interruption in bead production followed by sporadic ‘bursts’ of rapid droplet formation. At 1.6 % chitosan concentration, uniform droplets were observed, but the size was abnormally large (1454 ± 28 µm) – over 5 times the rectangular channel ducts width. Conversely, the solution at the lowest viscosity (1.2 % chitosan) generated beads with a diameter of 600±20 µm, which is roughly 2-3 times the width (**Figure 6A**.)

**Figure 6.**
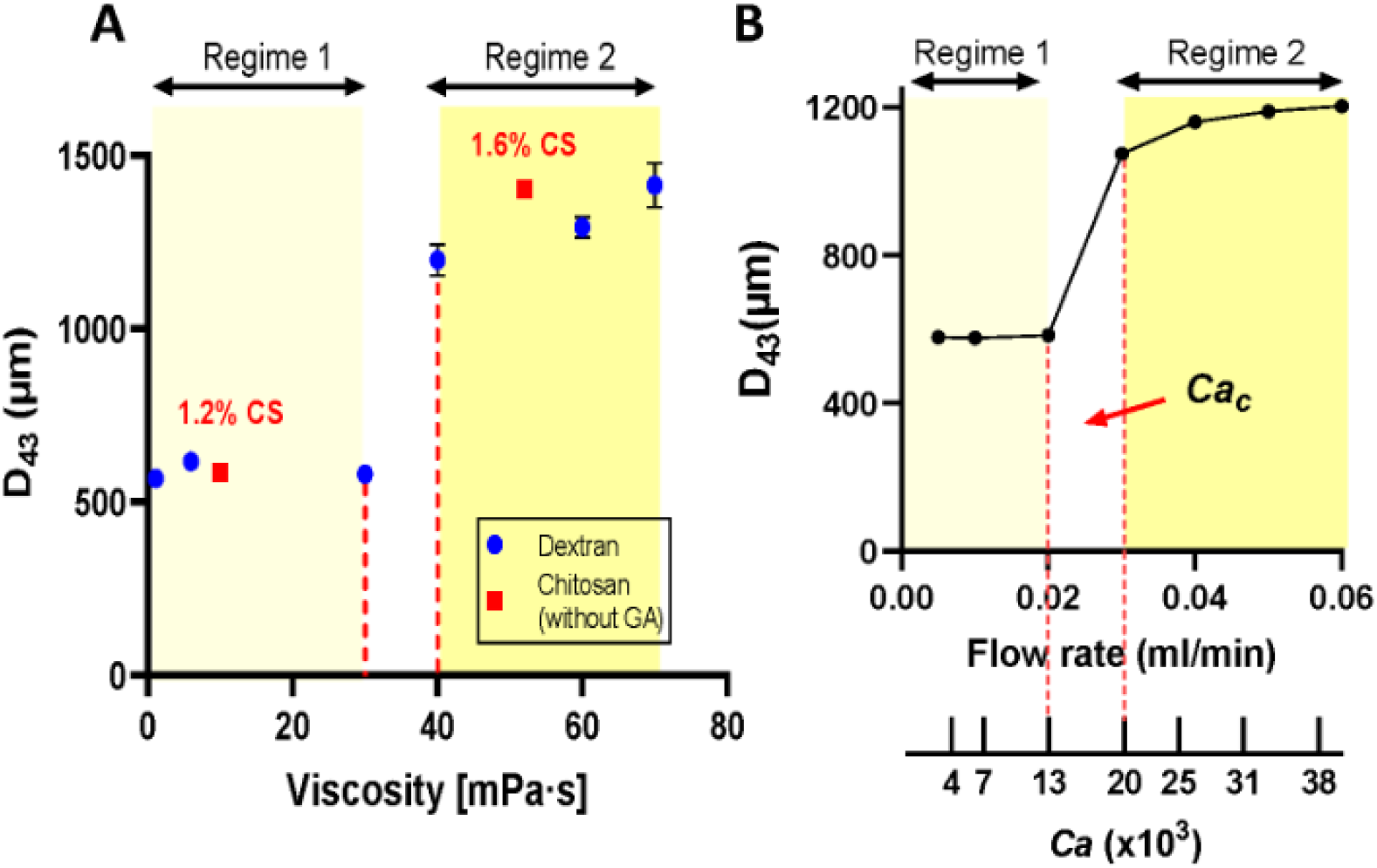
Effect of aqueous phase viscosity & flow rate on bead size. (**A**) Difference in average bead size depending on the aqueous phase viscosity at a fixed flow rate (0.1 mL/min). Dextran (blue dots) concentration was increased from 0 g/mL to 0.33 g/mL. The apparent viscosity was measured at shear rate ranging from 10-400 (1/s), which is calculated to be the shear rate range in the cylinder channel at flow rate (0.05-0.10 mL/min). (**B**) Effect of flow rate on bead size at a fixed chitosan concentration (1.6% w/v), (N = 3, n > 90). For both studies, Asym C was used. (*Ca* = Capillary number)

To further investigate the trend in bead size associated with the aqueous phase viscosity, we chose another biomaterial, dextran, which can dissolve in water at different concentrations to achieve different viscosities. In Figure 6.A, the *Regime 1* represents the region where the microchannel can consistently generate droplet sizes 2-3 times the rectangular duct width, whereas *Regime 2* represents the region with droplet size over 5 times the width. The 1.2 % chitosan solution falls under the *Regime 1* (Figure 6.A). The system allows the production of small droplets regardless of viscosity under a critical value, which was within 30-58 mPa∙ s in the current context. Although 1.2 % chitosan precursor solution allowed stable generation of small droplets, beads obtained when adding gelling agent did not withstand the rinsing process. Therefore, based on size uniformity and stability after the encapsulation process, 1.6 % chitosan precursor solutions were used for process optimization towards cell encapsulation.Flow transition associated with capillary number

To establish stable production of smaller microbeads, we dispersed the precursor solution (1.6 % w/v chitosan) at different flow rates and quantified the droplet diameter as a function of capillary number (*Ca*) ^23, 36^. The *Ca* was calculated using *Ca* = (*U*∙ƞ)/*γ* (*Ca* = capillary number, *U* = flow rate (m/s), ƞ = viscosity, and *γ* = interfacial tension, *γ* between the aqueous phase and the oil phase was assumed to be 35 mM/m). As viscosity changes as a function of time, the value used in *Ca* calculations was the viscosity of the chitosan precursor solution without the gelling agent, which was assumed to represent the viscosity immediately after mixing. Shown in **Figure 6B**, at flow rates below 0.02 mL/min (*Ca=* 1.3 × 10^−2^), the channel consistently generated small droplets (570 ± 50 µm) regardless of flow rate increase. When flow rates exceeded 0.03 mL/min (*Ca=* 2 × 10^−2^), the average diameter gradually increased as the flow rate increased. These observations suggest a transition between droplet formation mechanisms below vs above a critical *Ca* value (*Ca*_*c*_), with a transitional region between the two regimes.

This transition in droplet formation regimes is likely related to the balance between viscous force and interfacial tension ^37^. Below a critical *Ca* value (*Ca*_*c*_), which is observed to be within 1.3 × 10^−2^ to 2 × 10^−2^, interfacial tension mainly drives the droplet formation. However, both viscous force and interfacial tension influence the droplet generation above *Ca*_*c*_, resulting in flow transition where flow rate has more influence on bead diameter. Next, we calculated *Ca* of the results acquired from **Figure 6A** to determine if the values fall under the region correlated with the resulting bead diameter. The *Ca* for 1.2 % CS (at a flow rate of 0.1 ml/min) was calculated to be 0.01, which falls under *Regime 1* (below *Ca*_*c*_) and *Ca* for 1.6 % CS (at 0.1 min/ml) was 0.05, which was beyond *Ca*_*c*_ (*Regime 2*). The calculated results indicate that the same regime trend can be observed in chitosan precursor solution at different viscosity.

### Mechanical properties and injectability of microchannel vs stirred emulsification-generated beads

To determine whether the microchannel emulsification beads would be suitable for *in vivo* delivery, we compared their mechanical properties and resistance to passage through a needle to that of chitosan beads obtained through stirred emulsification. To allow side-by-side comparison, we generated stirred emulsification beads of matching volume-weighed bead diameter (*D[4,3]*) by adjusting the agitation rate during the process. We chose *D[4,3]* as metric since this is representative of the conditions the average MSC, which are distributed uniformly within the bead volume, would experience. The D[4,3] of microchannel emulsification beads was 765 ± 16 µm (SEM) with C.V of 9 ± 1 % (SEM) compared to the D[4,3] of stirred emulsification beads was 729 ± 50 µm (SEM) with C.V of 41 ± 5 % (SEM), as shown in **Figure 7**.

**Figure 7.**
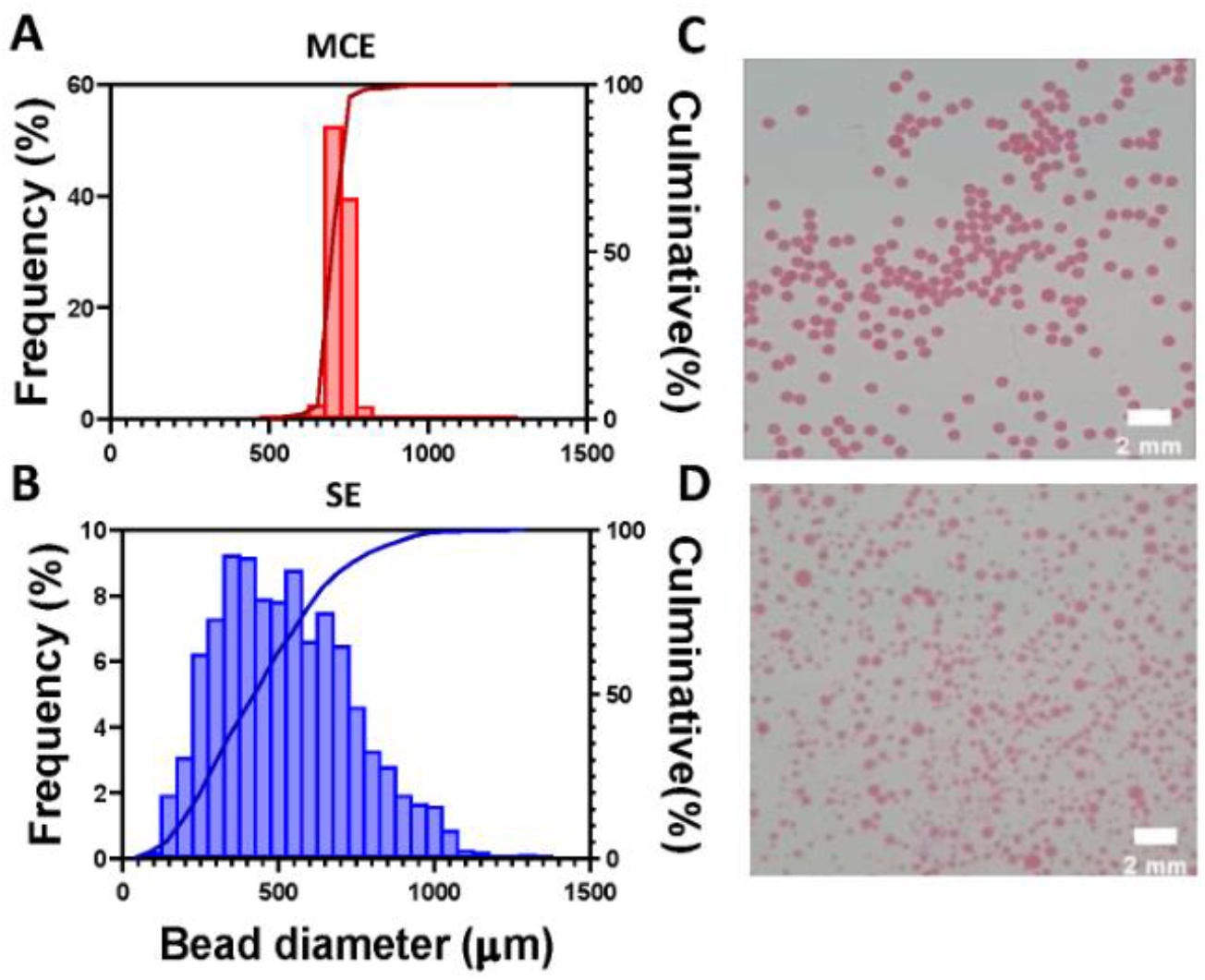
Size distribution of chitosan microbeads generated by (**A**) Microchannel emulsification or (**B**) Stirred emulsification, and the representative images of the respective beads (**C, D**). For the aqueous phase, 1.6% chitosan was used. To match the average size of microchannel emulsification beads, the stir speed was set to 600 rpm for SE. (N = 3, n > 750) MCE: microchannel emulsification. SE: stirred emulsificatio

To demonstrate improved injectability of beads with narrower size distribution, we infused the beads through a syringe needle that is slightly smaller in size (ID: 513 µm) compared to D[4,3]. To quantify the number of ruptured beads at different bead densities, we prepared beads suspended in two different volumes of HEPES buffered saline solutions. At both concentrations, the fraction of ruptured microchannel emulsification beads was significantly lower than that of stirred emulsification beads, namely 20 ± 5 % damaged beads/total beads compared to 39 ± 6 % at a high beads/HEPES volume ratio (1/3), reduced to 5 ± 5 % and 20 ± 3 % respectively at higher dilution (1/6) (**Figure 8A**). Reducing bead concentration may have significantly impacted the rheology of the solution passing through the needle when beads were uniform in size, but this effect may not have been as impactful for beads of broad size distribution. The fraction of ruptured beads is based on bead numbers, and hence the large fraction of ruptured beads in stirred emulsification would represent an even larger volumetric fraction of inadequately encapsulated cells. We also infused the beads through a smaller needle of 23 G (ID:337 µm) to measure the injectable limit. Most beads ruptured after injection for both microchannel emulsification and stirred emulsification beads, making it difficult to quantify.

**Figure 8.**
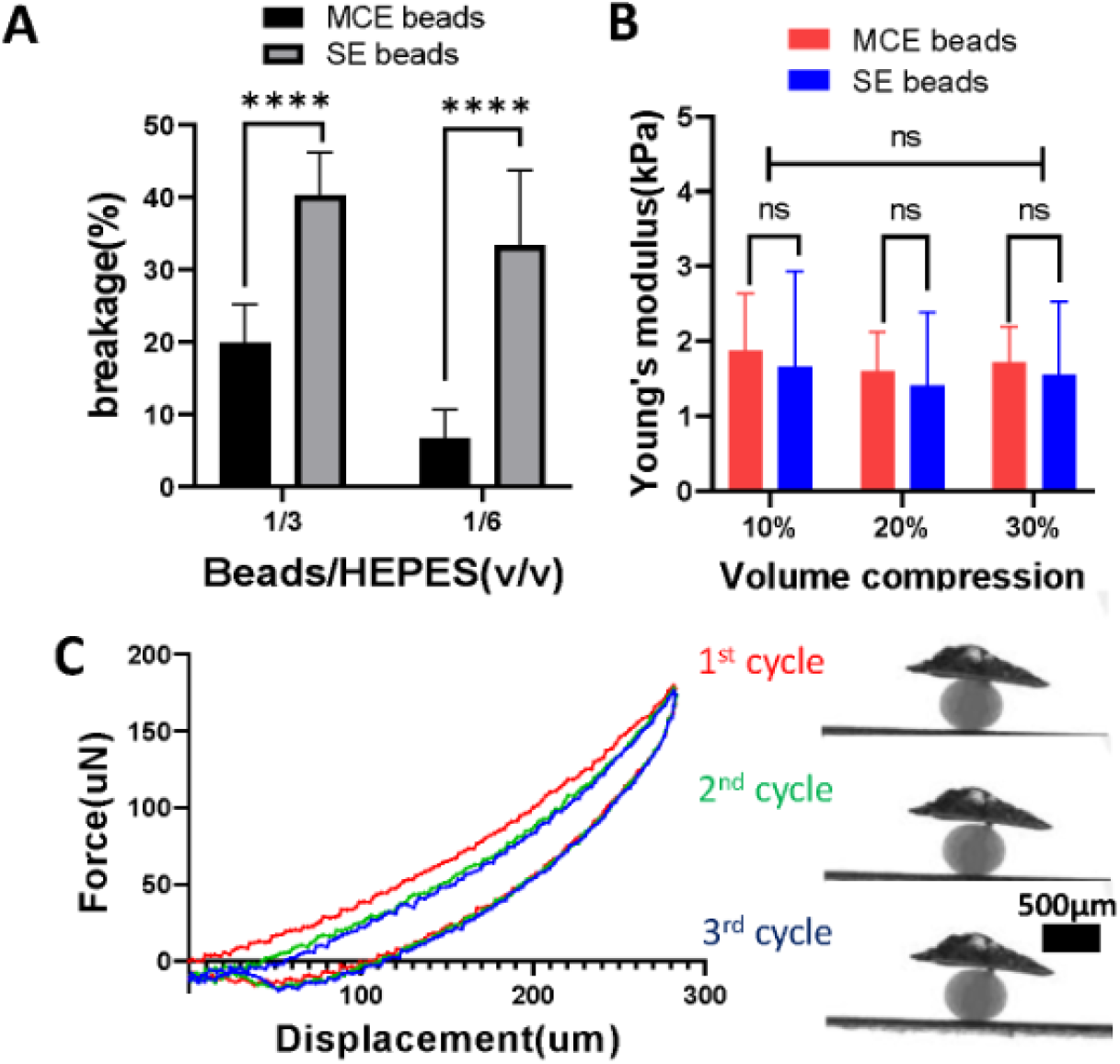
Mechanical properties of chitosan beads generate by microchannel versus stirred emulsification. (**A**) Percentage of beads ruptured after being injected through 21 G needle (ID: 513 µm). The beads were suspended in HEPES buffer in different dilution for injection (N=3, n > 20) (**B**) Compressive moduli of microbeads process by MCE and SE within 24 hours after production (N = 3, n = 6) (**C**) Representative viscoelastic behavior of microchannel emulsification beads compressed up to 30% volume for 3 cycles. MCE: microchannel emulsification. SE: stirred emulsification.

To further assess the mechanical properties of the beads, we used a parallel plate compression set-up to compress single beads of matching size (700-800 µm in diameter) up to 30 % in volume. The degree of volume compression was determined to match the inner diameter of the 21 G needle (513 µm) to the compressed bead height. No significant difference in compressive moduli was observed between microchannel emulsification and stirred emulsification beads, suggesting that the improved injectability of microchannel emulsification beads was resulting from the absence of undesirably large microbeads in the batch. After a first compression cycle, slight plastic deformation was observed as the force showed a negative value. Nonetheless, microchannel emulsification beads withstood 30 % volume compression for three consecutive cycles without any rupture (**Figure 8C**).

### Viability of immobilized MSCs and VEGF secretion

Next, we measured the viability of MSCs after the bead production process to assess (a) the suitability of the process for cell immobilization and (b) the encapsulated cells’ ability to secrete VEGF A, the main growth factor involved in neovascularization^38^, in the low serum mimicking the condition observed in ischemic tissues. MSCs showed high viability in both microchannel and stirred emulsification processes 1 hour after encapsulation (Day 0), as shown in **Figure 9A, B**. To reflect conditions required for eventual cytokine secretion and transplantation assays, the cells were incubated in alpha-MEM with FBS 0.2% v/v (low serum) for 3 days. As expected in low-serum conditions ^39^, the viability decreased over the following 3 days but no significant differences in cell survival nor VEGF secretion were observed between the microchannel and stirred emulsification beads (Figure 9C). These results offer preliminary evidence that the microchannel emulsification process would be a suitable alternative to stirred emulsification for MSC encapsulation and delivery for ischemic tissue repair.

**Figure 9.**
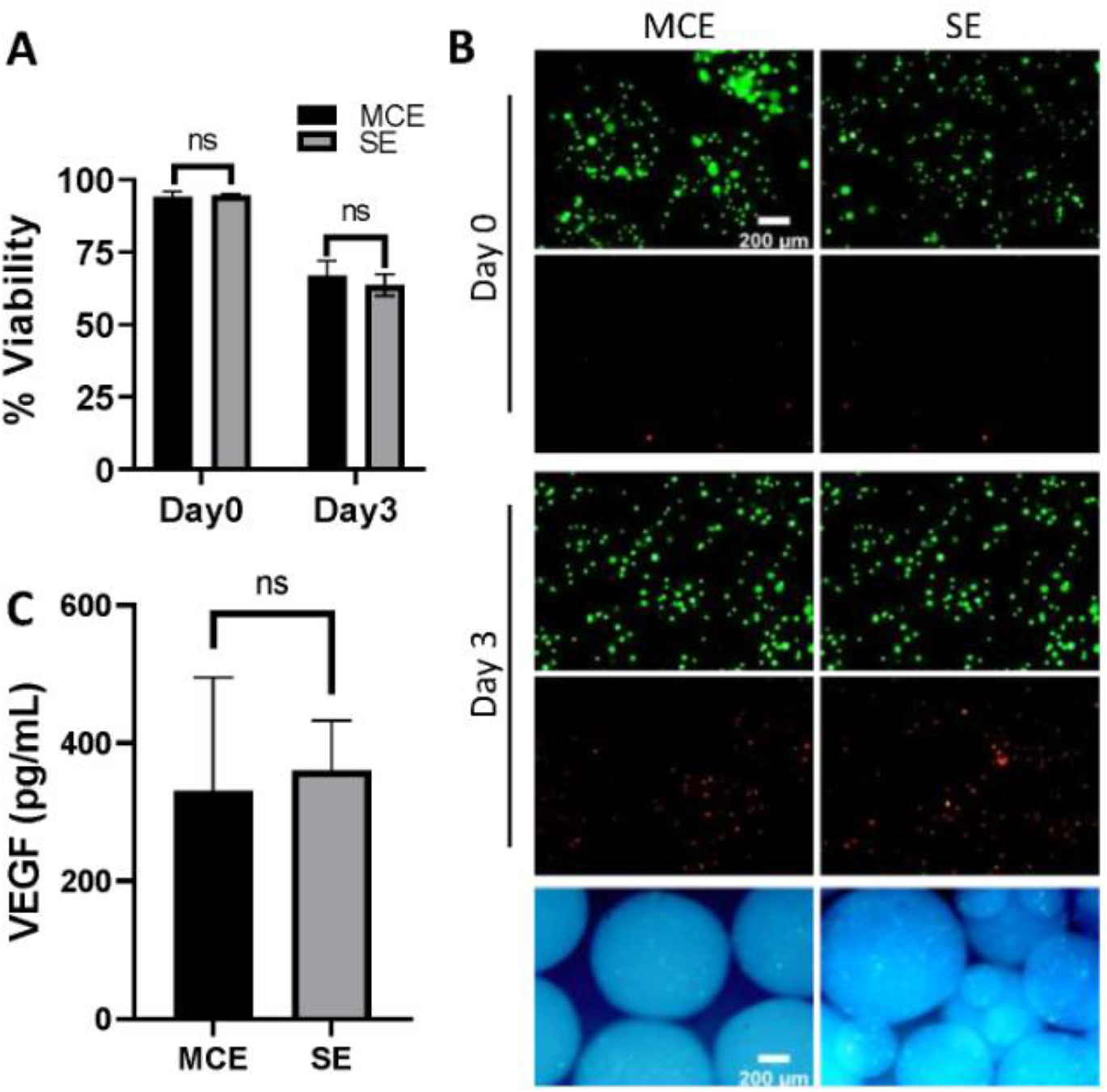
Cytocompatibility of microchannel and stirred emulsification beads. (**A**) Quantified viability of hMSCs immobilized in microbeads via each process (N=3). (**B**) Representative fluorescence images of immobilized hMSCs. (**C**) VEGF release in culture media after 3 days of culture in chitosan microbeads (N=3). MCE: microchannel emulsification. SE: stirred emulsification.

### Perspectives on adapting microchannel emulsification for chitosan-based single cell encapsulation

In our previous study ^23^, we designed a prototype microchannel emulsification device to immobilize pancreatic cell aggregates (MIN6 cells) in alginate microbeads. Here, both the cell type and gelation mechanisms were modified requiring adaptation of the microchannel emulsification process. Initially, the same channel geometry (symmetric channels) was used with chitosan as previously applied for alginate bead production. However, contrary to the alginate-based encapsulation process, where the droplets gel immediately after being detached from the channel, bead coalescence was observed since heat-induced gelation of chitosan is not instantaneous. To prevent droplet coalescence and to reach the target bead diameter (~600 µm to allow needle injection), we added fluorosurfactant and reduced the bead size significantly using asymmetric channels. It was assumed that the cylinder shaft of the asymmetric channel limited the volume of the aqueous phase entering into the channel, while the large aspect ratio of the slot allowed the oil to fill the slot prior to droplet formation and constricted the droplet neck as the aqueous phase exited the shaft ^35^.

Due to the dimension tapering during laser micromachining, it was difficult to produce a channel size (inlet cylindrical diameter) smaller than 220 µm for asymmetric channels. As the channel size decreased, smaller droplets formed which accelerated heat transfer and hence gelation. This improved bead stability in the collection compartment, but also led to increased time-dependent changes in viscosity affecting the dependence of bead size and process stability on flow rate. To improve process robustness, we studied the effect of aqueous phase viscosity and flow rate, and hence *Ca*, on bead size and process stability (uninterrupted bead production). Since the viscosity of the chitosan precursor solution changed over time during dispersion, it was crucial to determine *Ca*_*c*_ and make necessary changes in either the viscosity or the flow rate.

The device also went through structural changes (Supporting information). The volume of aqueous phase required to fill the bottom chamber before dispersion through a channel was reduced from 9 mL to 1.5 mL to minimize the aqueous phase dead volume. The second outlet in the bottom chamber previously designed to prevent pressure build-up was removed as the pressure loss through the outlet could cause bead production instability. To reduce the probability of early bead-bead contact in the collection vessel, the surface area of the collection site was increased from 60 cm^2^ to 120 cm^2^.By using a fluorosurfactant, adapting channel and device geometry, as well as changing process variables, we were able to produce uniform-size chitosan beads (C.V < 10%) of diameters below 600 µm without incurring coalescence. When cells were added in the precursor solution, the bead size obtained increased to ~750 µm likely due to increases in viscosity associated with the presence of cells. We observed intermittent channel obstruction or production of abnormally large droplets 30 min after the initial aqueous phase dispersion during the cell encapsulation. To maintain the size uniformity of a batch, the production time was limited to ensure that the viscosity change during the process did not reach values leading to changes in droplet formation regime. While maintaining improved size uniformity and injectability, no significant difference in cell viability and VEGF secretion was observed compared to the previous method (stirred emulsification).

### Impact and future directions

The objective of this research was to develop and optimize a microchannel emulsification method for the fabrication of chitosan hydrogel microbeads with control over process stability, bead size, bead mechanical properties, and viability of encapsulated cells. The microchannel emulsification device was successfully adapted for the thermoresponsive chitosan-based gelation process by redesigning the device and providing a temperature gradient in the oil phase. Coalescence of smaller (˂1500 µm diameter) beads was prevented by adding surfactants, allowing production of chitosan beads of uniform size with high MSC survival after the process.

A significant advantage of microchannel emulsification over stirred emulsification-based encapsulation of MSCs for cell therapy applications is size uniformity. This was demonstrated through the coefficient of variation (<10% versus > 40% respectively) and through the reduction in the fraction of damaged beads after passage through a 21 G needle, likely due to a smaller number of beads with diameter largely exceeding the syringe inner diameter. The reduction in bead rupture was not attributed to differences in compressive modulus, as no significant differences between microchannel emulsification and stirred emulsification beads were observed for this metric. Size uniformity is important to avoid broken beads but also to ensure that all injected cells behave similarly and that the cell payload and biological effect of the cell therapy is well controlled and reproducible.

The microchannel emulsification device currently can adopt alginate-based and chitosan-based gelation processes. The system’s versatility would likely allow adaptation to other gelation processes and polymers for various biopharmaceutical, food and drug encapsulation applications ^40^.

Obtaining smaller beads may be desirable for future clinical applications in the context of cell-based therapies. Although the bead diameter was significantly reduced using asymmetric channels, decreasing the bead size to 400 µm in diameter is desired as smaller needles (23-25 G) are commonly used for subcutaneous injections ^22^. This can be done by using a laser setup that can minimize dimension tapering to generate channels with smaller slot widths (< 100 µm). One solution would be to develop a technique that allows the laser system to mill the channel from both sides of the PTFE plate. In parallel, the aqueous phase viscosity and flow rate should be further adjusted to let the system *Ca* fall under the stable processing zone. Integrating temperature-controlled zones within the device could afford better control over chitosan gelation kinetics.

To achieve higher flow rates and reduce viscosity changes during processing, multiple channels can be created in a single PTFE plate to promote simultaneous bead production at each channel. The system should also be optimized to establish a stable bead production at higher cell density, considering that many clinical applications require cell density over 10^6^ cells/mL hydrogel to promote tissue regeneration ^39, 41^. These efforts should account for the viscosity rise and changes in *Ca* values resulting from the cell density increase. The effect of different chitosan formulations on the MSC secretome and *in vivo* retention at the delivery site should be investigated to maximize their immunomodulatory, angiogenic and other therapeutic effects.

## CONCLUSIONS

MSCs have shown great potential as regenerative medicine for their immunomodulatory functions and secretion of pro-angiogenic cytokines. To ensure their long-term survival and maximize their therapeutic effect, the cells can be immobilized in microbeads to offer better retention and mechanical and/or immune protection while allowing a bidirectional diffusion of nutrients, oxygen, and biological signals. For applications requiring cell release from the microbeads, bioresorbable polymers such as chitosan are preferred over alginate which is not readily degraded by endogenous human enzymes. We previously described a stirred emulsification approach to encapsulate MSCs in chitosan microbeads, but the broad bead size distribution posed a challenge in maximizing cell viability and injectability. In this work, a microchannel emulsification-based process was adapted for MSC encapsulation in a thermoresponsive chitosan hydrogel. Process parameters were adjusted to generate monodisperse chitosan beads of with controlled sizes ranging from 600 µm – 1500 µm in diameter while maintaining high MSCs viability (95 %) after the process. Microchannel emulsification is an attractive option to produce uniform-size chitosan and other thermoresponsive polymer-based microbeads for cell therapy, drug delivery, and food applications.

## ACKNOWLEDGEMENTS

We thank Balaji Ramachandran and Jonathan Brassard for valuable discussions, and Dr. Ramachandran also conducted interfacial tension measurements with Dongjin Shin. This study was funded by the Quebec Cell, Tissue and Gene Therapy Network –ThéCell (a thematic network supported by the Fonds de recherche du Québec–Santé). The work was also supported by grants from the Canada Foundation for Innovation (35507), and the Natural Sciences and Engineering Research Council of Canada (CAH: RGPIN-2020-05877 and SL: RGPIN-2020-06684). This research was undertaken, in part, thanks to funding from the Canada Research Chairs Program: CAH holds the Canada Research Chairs in Cellular Therapy Bioprocess Engineering. DSS was supported by Eugenie Ulmer-Lamothe awards from the Department of Chemical Engineering, McGill University.

## Notes

### Competing Interest Statement

The authors have declared no competing interest.

