## Supporting Information for "Mesenchymal stromal cell encapsulation in uniform chitosan beads using microchannel emulsification"

### Supplementary equations: Modified Hertzian half space contact model was used to calculate the compressive modulus of chitosan microbeads.

$a=\left( R-\delta\right)\tan\left( \varphi\right)$	(Equation S1)

$f\left( a \right)= \frac{2\left( 1+\nu\right)R^{2}}{{(a^{2}+4R^{2})}^{3/2}} + \frac{1-\nu^{2}}{{(a^{2}+4R^{2})}^{1/2}}$	(Equation S2)

$E= \frac{3\left( 1-\nu^{2} \right)F}{4\delta a} - \frac{f\left( a \right)F}{\pi\delta}$	(Equation S3)

where $E$= compressive modulus $F$ = applied force during the loading time in the first compression cycle, δ = displacement, *a* = radius of the contact area, $\varphi$= contact angle, *R* = microbead radius, *v* = Poisson’s ratio was assumed to be 0.5.

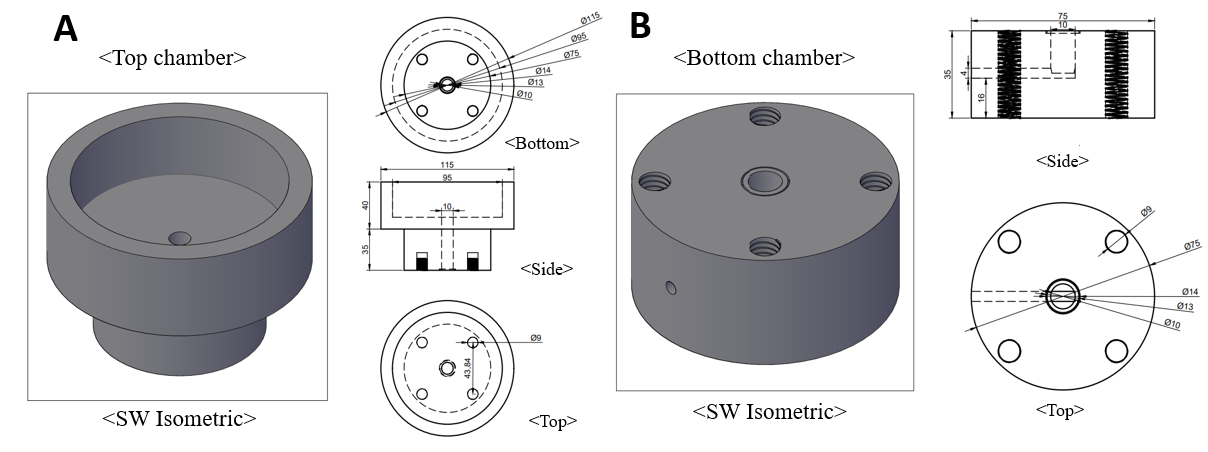

**Figure S1.** Computer-assisted design of the microchannel emulsification device. **(A)** Top chamber CAD design, **(B)** Bottom chamber CAD design. Four threaded holes (9 mm in diameter) were machined through the bottom chamber and the top chamber. Two Face seals type O-ring grooves were machined on both chambers. Two O-rings (5233T297, ID: 12 mm, OD: 14 mm, McMaster-Carr, CA) were fitted into both chambers to press down the microchannel plate from both sides.

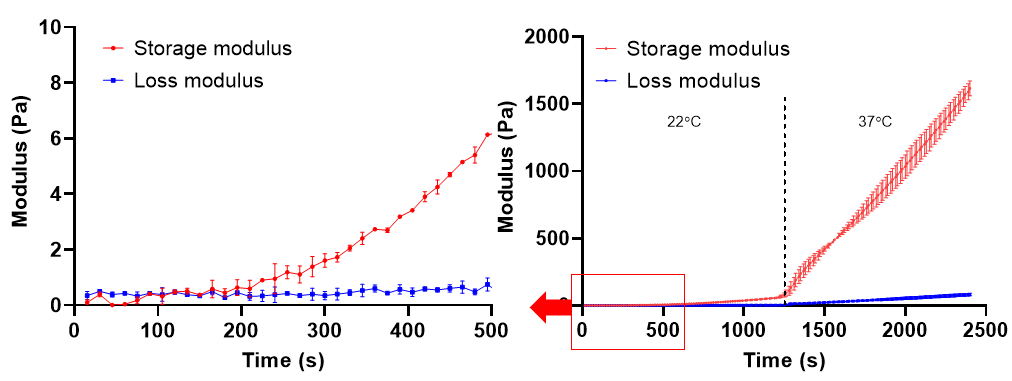

**Figure S2.** Storage modulus (G’) and Loss modulus (G’’) change in time immediately after mixing chitosan solution (1.6% w/v), gelling agent (PB 0.02M/SHC 0.075M), and cell culture media (10% BSA). The solution was first kept at 22°C for 20 min, and then temperature was increased to 37 °C to mimic the behavior in the heated oil phase (MCR301 rheometer, CC10 geometry, 1 Hz, 1% strain) (mean ± SD, N=3).
